# Structural insights of a highly potent pan-neutralizing SARS-CoV-2 human monoclonal antibody

**DOI:** 10.1101/2021.09.28.462234

**Authors:** Jonathan L. Torres, Gabriel Ozorowski, Emanuele Andreano, Hejun Liu, Jeffrey Copps, Giulia Piccini, Lorena Donnici, Matteo Conti, Cyril Planchais, Delphine Planas, Noemi Manganaro, Elisa Pantano, Ida Paciello, Piero Pileri, Timothée Bruel, Emanuele Montomoli, Hugo Mouquet, Olivier Schwartz, Claudia Sala, Raffaele De Francesco, Ian A. Wilson, Rino Rappuoli, Andrew B. Ward

**Author notes:** These authors contributed equally to this work (Jonathan L. Torres, Gabriel Ozorowski).

## Abstract

As the coronavirus disease 2019 (COVID-19) pandemic continues, there is a strong need for highly potent monoclonal antibodies (mAbs) that are resistant against severe acute respiratory syndrome-coronavirus 2 (SARS-CoV-2) variants of concern (VoCs). Here, we evaluate the potency of a previously described mAb J08 against these variants using cell-based assays and delve into the molecular details of the binding interaction using cryo-EM. We show that mAb J08 has low nanomolar affinity against VoCs, binds high on the receptor binding domain (RBD) ridge and is therefore unaffected by most mutations, and can bind in the RBD-up and -down conformations. These findings further validate the phase II/III human clinical trial underway using mAb J08 as a monoclonal therapy.

**One Sentence Summary:** Potent neutralizing monoclonal antibody J08 binds SARS-CoV-2 spike independent of known escape mutations.

## Main Text

To date, over 219 million cases and 4.5 million deaths worldwide have been caused by SARS-CoV-2 (*1*), along with high levels of unemployment and far-reaching supply chain issues in the world’s economy. While the situation is vastly improving with numerous vaccine roll-outs and over 6.03 billion doses of vaccines administered worldwide, these have been heavily skewed towards developed nations (*2*). Hence, disadvantaged communities and low- and middle-income countries remain potential hotspots for emergence of viral variants with increased infectivity and mortality. Designated variants of concern (VoCs) pose the greatest threat to progress made thus far because data have shown that they can reduce the effectiveness of the COVID-19 vaccines (*3*). These findings have led vaccine manufacturers to test booster shots (*4*–*6*) that will ultimately result another logistical distribution and administration challenge.

COVID-19 vaccines, whether mRNA, protein, or viral vector based, aim to provide acquired immunity and protection against serious disease by presenting the SARS-CoV-2 spike protein (S-protein) to the immune system (*7*). The S-protein is a glycosylated, homotrimeric type I transmembrane fusion protein responsible for host cell attachment via its receptor binding domain (RBD) to human angiotensin-converting enzyme 2 (hACE2), a process which is enhanced by a co-receptor, cellular heparan sulfate (*8*). The S-protein is made up of the S1 domain (residues 1 to 685) containing the RBD and N-terminal domain (NTD) and the S2 domain (residues 686 to 1213) housing the fusion machinery. Both the RBD and NTD are immunodominant epitopes that are targeted by a majority of neutralizing antibodies (*9*).

Ending the pandemic is being challenged by the emergence of VoCs, uneven roll-out of vaccines worldwide, vaccine hesitancy, uncertainty of whether COVID-19 vaccines can prevent transmission, breakthrough infections amongst vaccinated populations, and if vaccinated and naturally infected individuals will have long-lasting immunity against the virus. Therefore, having readily available, highly potent, and viral variant resistant monoclonal antibodies may serve to mitigate the propagation of troublesome variants worldwide and treat those that remain vulnerable after vaccination. Early efforts with monoclonal therapies have had mixed results. LY-CoV555 (bamlanivimab), developed by Eli Lilly and Company, was found to have no significant effect on viral load compared to a placebo in phase II trials, and is easily susceptible to escape mutations present in common variants of concern (*10*, *11*). Monoclonal cocktails are one strategy to decrease the chance of viral escape, and a recent Emergency Use Authorization was given for Regeneron’s casirivimab (REGN10933) and imdevimab (REGN10987), and Eli Lilly and Company’s bamlanivimab and etesevimab combinations (*10*, *12*).

In this study, we characterized the neutralization breadth of a highly potent human monoclonal antibody, J08, previously isolated from a convalescent COVID-19 patient (*13*), against the SARS-CoV-2 B.1.1.7 (Alpha), B.1.351 (Beta), P.1 (Gamma) and B.1.617.2 (Delta) VoC. Following functional characterization, we further investigated the structural details of J08 to define the epitope that retains neutralization activity against all VoCs.

### J08 cross-neutralizes all current SARS-CoV-2 variants of concern

To evaluate neutralization breadth, J08 was tested for binding, ACE2 blocking, and neutralization against the SARS-CoV-2 D614G virus and VoCs B.1.1.7 (isolated in the United Kingdom) (*14*), B.1.351 (isolated in South Africa) (*15*), P.1 (isolated in Brazil) (*16*), and B.1.617.2 (isolated in India) (*17*). These VoCs have been re-named by the World Health Organization (WHO) as Alpha, Beta, Gamma, and Delta variants, respectively (*18*). We first tested antibody binding or ACE2 blocking in an enzyme-linked immunosorbent assay (ELISA) using the S-protein RBD from the original Wuhan strain and from the Alpha through Delta VoCs. J08 was able to bind and interfere with the RBD/ACE2 interaction with all tested variants (fig. 1, A and B). We next evaluated the neutralization activity of J08 against authentic SARS-CoV-2 and VoC viruses using a cytopathic effect-based microneutralization (CPE-MN) assay, an S-2 fuse neutralization assay, and a SARS-CoV-2 pseudovirus platform. The neutralization experiments were performed in three different and independent laboratories. The CPE-MN assay demonstrated that J08 was able to neutralize the SARS-CoV-2 D614G virus with a 100% inhibitory concentration (IC_100_) of 3.9 ng/mL and maintain its extremely high neutralization activity against all tested VoCs with an IC_100_ of 3.9, 9.7, 4.9 and 6.2 ng/mL for the Alpha, Beta, Gamma, and Delta VoCs, respectively (fig. 1, C and F). A similar scenario was observed with the S-fusion neutralization assay, where J08 was able to neutralize all VoCs tested. Given the extremely high neutralization potency of J08, we were not able to define a 50% inhibitory concentration (IC_50_) against the D614G, Alpha, and Delta variants and therefore we assigned it as <1 ng/mL. On the other hand, it was possible to define the neutralization potency of J08 against the Beta and Gamma variants that showed an IC_50_ of around 3.2 and 1.0 ng/mL, respectively (fig. 1, D and F). Finally, we evaluated the neutralization activity of J08 using a lentiviral pseudovirus platform produced with a SARS-CoV-2 spike variant, which included a cytoplasmic tail deletion of 19 amino acids. Similar cytoplasmic tail deletions showed an enhanced spike incorporation into pseudovirions and increased viral entry into cells, compared to those with full-length S-protein (*19*). With our pseudovirus platform, we observed an overall lower neutralization IC_50_ by J08. The differences in measured inhibitory concentrations between assay types could arise from the pseudovirus platform used and its high incorporation of S-protein into pseudovirions, as well as the use of cell lines showing different levels of cell surface hACE2 presentation. Nonetheless, J08 showed an IC_50_ against the D614G, Alpha, Beta, Gamma, and Delta variants of 22, 77, 499, 147 and 226 ng/mL, respectively, further confirming its ability to neutralize all VoCs (fig. 1, E and F).

**Fig. 1.**
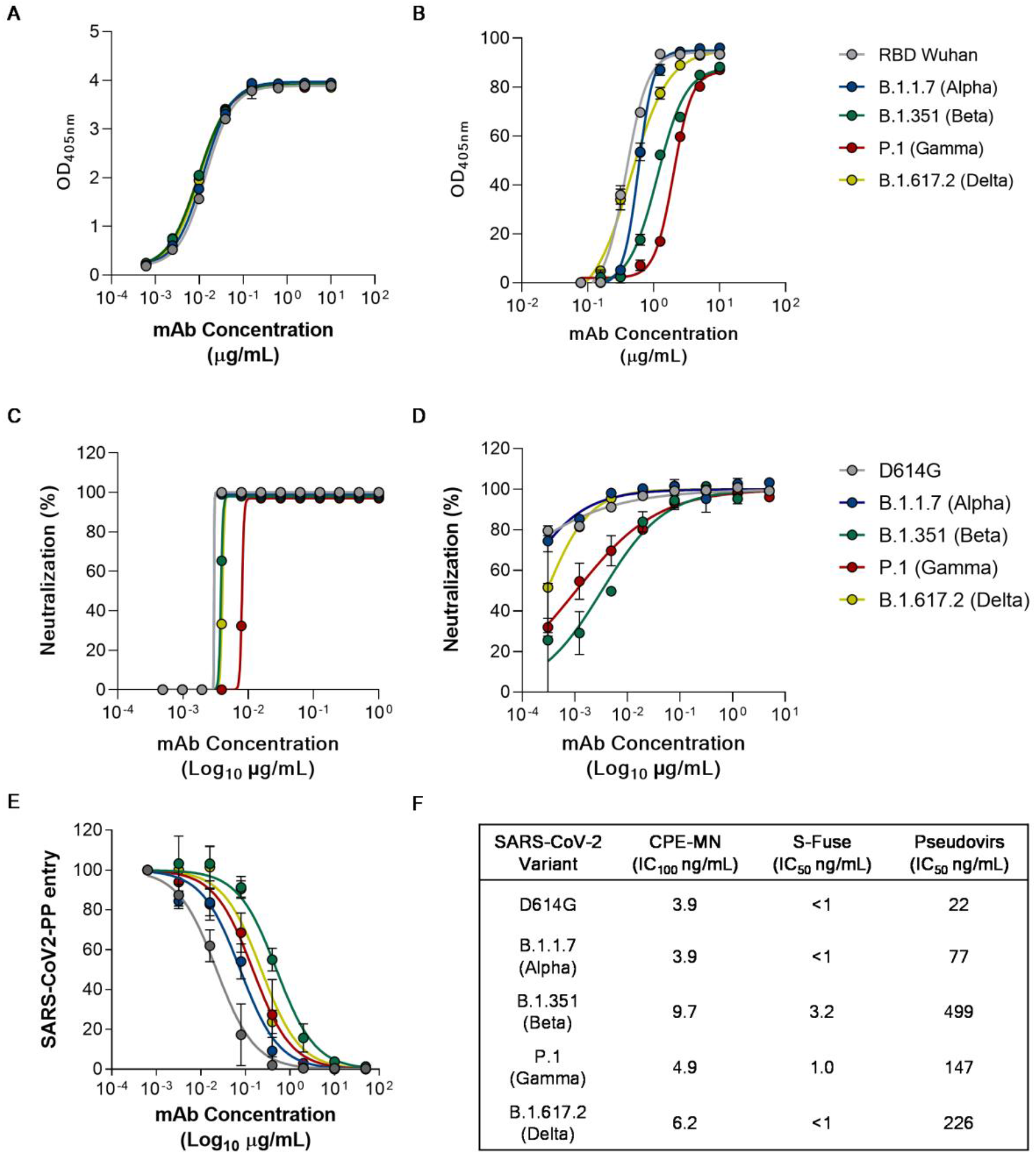
J08 activity against SARS-CoV-2 and emerging variants. Graphs show the ability of mAb J08 to bind (A), block RBD/ACE2 interaction (B), and neutralize SARS-CoV-2 D614G, B.1.1.7, B.1.351, P.1 and B.1.617.2 using a CPE-MN (C), S-fuse (D) and pseudovirus platform (E). (F) The table summarizes the IC_100_ and IC_50_ results obtained for all neutralization assays.

### Neutralization activity of competitor antibodies against SARS-CoV-2 and emerging variants

To compare the neutralization potency and breadth of J08 to other SARS-CoV-2 neutralizing mAbs that have received emergency use authorization for COVID-19 treatment, we recombinantly expressed REGN10987 (*12*), REGN10933 (*12*), S309 (*20*), CoV2.2196 (*21*), and LY-CoV016 (*10*) as immunoglobulin G1 (IgG1). All mAbs were tested by the CPE-MN assay using authentic SARS-CoV-2 viruses and by our pseudovirus platform against the D614G, Alpha, Beta, Gamma, and Delta VoCs. REGN10987 showed a neutralization potency of 24.6, 19.5, 4.9, 3.1 and 19.7 ng/mL against D614G, Alpha, Beta, Gamma and Delta VoCs, respectively (fig. S1, A and K). A similar trend for these variants was observed with our pseudovirus platform, although the Delta variant showed a 52-fold reduction (fig. S1, F and K). Antibody REGN10933, which recognizes the receptor binding motif of the spike protein, showed high neutralization potency against D614G, Alpha, and Delta VoCs, but was heavily impacted by the Beta and Gamma VoCs, showing a 59 and 29.5-fold decrease respectively (fig. S1, B and K). Our pseudovirus platform results were in accordance with the CPE-MN assay (fig. S1, G and K). We then evaluated antibody S309, which targets a region outside of the S-protein receptor binding motif (*20*). This antibody, when assessed by the CPE-MN assay, retained its neutralization potency against all SARS-CoV-2 variants showing an IC_100_ of 156, 248, 78, 25 and 79 ng/mL for the D614G, Alpha, Beta, Gamma and Delta VoC, respectively (fig. S1, C and K). In our pseudovirus platform, S309 was able to neutralize all SARS-CoV-2 variants but in the 4 – 41 μg/mL range (fig S2, H and K). Antibody CoV2-2196 showed high neutralization potency against all variants in both the CPE-MN assay and pseudovirus platform ranging from 12.3 – 49.2 ng/mL and 45.0 – 425.0 ng/mL, respectively (fig. S1, D, I and K). Despite CoV2-2196 being the only antibody that showed high neutralization potency against all variants, J08 remains the only antibody able to neutralize all VoC with an IC_100_ below 10 ng/mL. Finally, we evaluated antibody LY-CoV016 which showed to be heavily impacted by the Alpha variant with a 12-fold decrease in neutralization and unable to bind to the Beta and Gamma variants (fig. S1, E and K). LY-CoV016 lost activity to Alpha, Beta and Gamma, but was not impacted by the Delta VoC showing an IC_100_ of 49.6 ng/mL (fig. S1, E and K). When LY-CoV016 was tested in our pseudovirus platform, it was able to neutralize the D614G and Delta variants with a similar potency of around 300 ng/mL (fig. S1, J and K). These results are consistent with previously reported neutralization activity of each antibody (*11*, *22*–*27*).

### J08 can bind to RBD -up and -down conformations

For structural studies, we generated two constructs based on SARS-CoV-2-6P (six proline): one with the RBDs restricted to the down configuration by introduction of an interprotomer disulfide bond at positions C383 and C985 (Mut2) (*28*), and a second with unrestricted RBD movement but higher stability, containing an interprotomer disulfide bond at C705 and C883 (Mut7). Single particle cryo-EM analysis of two complexes, SARS-CoV-2-6P-Mut2 + Fab J08 and SARS-CoV-2-6P-Mut7 + Fab J08, resulted in 4 cryo-EM maps and associated models: SARS-CoV-2-6P-Mut2 trimer alone (3.2 Å), SARS-CoV-2-6P-Mut2 + Fab J08 Conformation 1 (3.4 Å), SARS-CoV-2-6P-Mut2 + Fab J08 Conformation 2 (3.4 Å), and SARS-CoV-2-6P-Mut7 + Fab J08 Conformation 3 (4.0 Å) (fig. 2A; fig. S2; table S1). Although Fab J08 has nanomolar affinity to the trimer, the short incubation time during sample preparation for cryo-EM enabled reconstruction of a trimer with no antibody bound, serving as a comparator for our analysis with J08-bound structures. Lastly, we determined a 2.54 Å crystal structure of recombinant RBD in complex with Fab J08 to address details in the antibody-antigen interface (table S2).

**Fig. 2.**
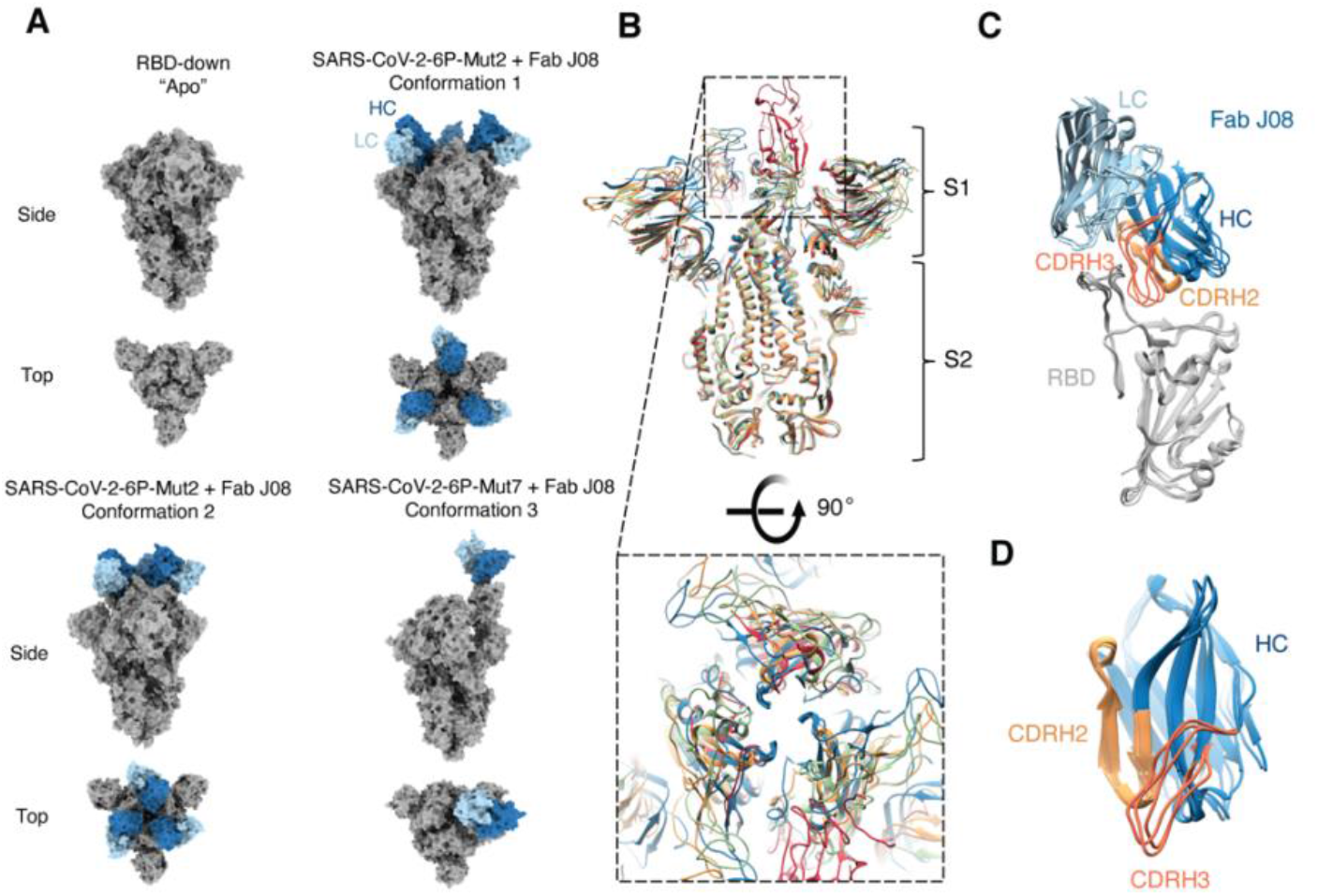
Conformational states of the S-protein-Fab J08 complex observed by Cryo-EM. (**A**) Top and side view surface representations of SARS-CoV-2-6P + Mut2, SARS-CoV-2-6P-Mut2 + Fab J08 Conformation 1, SARS-CoV-2-6P-Mut2 + Fab J08 Conformation 2, and SARS-CoV-2-6P-Mut7 + Fab J08 Conformation 3. S-protein is labeled in gray, heavy chain (HC) in dark blue, and light chain (LC) in light blue. (**B**) Side and top views of a slice through section of superimposed models of SARS-CoV-2-6P-Mut 2, SARS-CoV-2-6P-Mut 2 + FabJ08 (Conformations 1 and 2), and SARS-CoV-2-6P-Mut 7 + Fab J08 (Conformation 3) reveals flexibility at the S1 domain which affects the opening at the apex. (**C**) The structure of the RBD (gray) across Conformations 1-3 does not change when Fab J08 is bound. (**D**) CDRH3 (salmon) of Fab J08 exhibits more movement to accommodate binding to the RBD -up or -down conformations. On the other hand, CDRH2 (light orange) shows less movement across the different models.

In the SARS-CoV-2-6P-Mut2 + Fab J08 complex, we captured two different poses of Fab J08 through 3D classification, which we call Conformation 1 and Conformation 2. In Conformation 1, the J08 Fabs are further apart with a more closed apex (RBD more down), while Conformation 2 has the Fabs closer together and a more open apex (RBD slightly open) (fig. 2A). Superimposition of the S-protein models revealed movement in the S1 subunit, in comparison to minimal movement in the S2 subunit (fig. 2B). Both conformations consist of 3 Fab molecules bound to a single S-protein trimer. In the SARS-CoV-2-6P-Mut7 + Fab J08 complex (Conformation 3), Fab J08 bound to one RBD-up while the two other RBDs were down. Unlike the Mut2 complexes, most particles had a stoichiometry of only 1 Fab per S-protein trimer (figure 2A).

Alignment of the RBD-antibody portion of each of the models revealed no major differences in the angle of the Fab relative to the RBD, and the epitope remained constant (fig. 2C). However, more subtle differences were observed within the epitope-paratope interaction. The heavy chain complementarity-determining region 2 (CDRH2) loops were rigid, while the heavy chain CDRH3 loops were more variable, suggesting that CDRH2 was the anchoring interaction while the CDRH3 was slightly more plastic and could accommodate either the RBD-down or -up conformation (fig. 2D). These subtle differences might also be a result of differences in resolution across the various datasets. Concomitant with these structural differences, we observed variable buried surface areas (BSA) for Fab J08 bound to the 3 different conformations. Although J08 binds to the same epitope across the different conformations we observed that Fab J08 has the smallest footprint on the RBD in Conformation 3 at 658 Å^2^, followed by Conformation 1 at 728 Å^2^, and Conformation 2 at 985 Å^2^ (fig. 3A). When not under the constraints of an intact protomer or trimer, the BSA between J08 and RBD based on the x-ray structure is 676 Å^2^, most similar to Conformation 3.

**Fig. 3.**
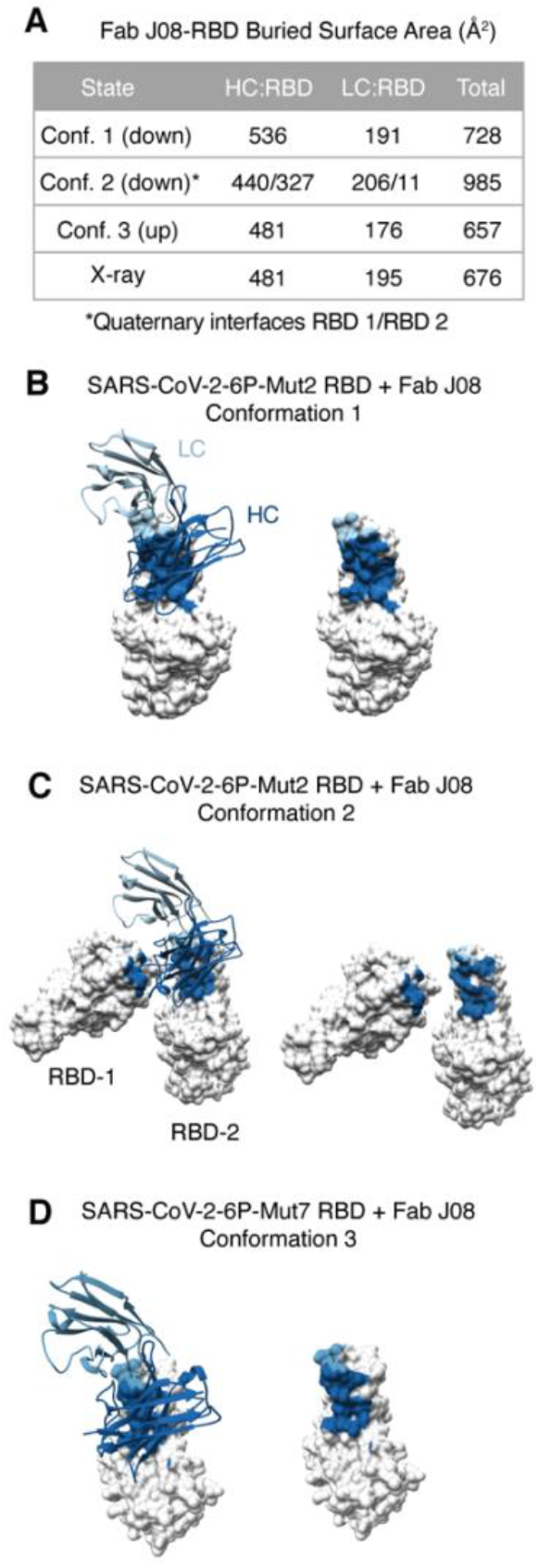
BSA and epitope footprint of the three conformations of the S-protein + Fab J08 complex. (**A**). Conformation 3 has the smallest calculated BSA (657 Å^2^), followed by the x-ray structure (676 Å^2^), Conformation 1 (728 Å^2^), and finally Conformation 2 (986 Å^2^). Surface representation of the RBD (gray) in complex with Fab J08 (HC dark blue, LC light blue) for SARS-CoV-2-6P-Mut2 RBD + Fab J08 Conformation 1 (**B**), SARS-CoV-2-6P-Mut2 RBD + Fab J08 Conformation 2 (**C**), and SARS-CoV-2-6P-Mut7 + Fab J08 Conformation 3 (**D**).

J08 can accommodate the dynamic movement of the RBD in the down or up position with several key contacts (fig. 3, B to D; table S3). CDRH2 appears to play an anchoring role and provides the most hydrogen bonds (including salt bridges) between antibody and RBD in Conformations 1 and 2. In the -up Conformation 3, the antibody is tilted slightly increasing the number of CDRH3 contacts, consistent with the x-ray structure of the RBD-Fab J08 complex (table S3). Accordingly, in Conformations 1, 3, and the x-ray structure, the heavy chain interface residues (defined as contributing greater than 5 Å^2^ BSA) all reside in CDRH2 and CDRH3 (tables S3 and S4). A portion of the interprotomer Fab contacts in Conformation 2 are provided by CDRH1 (tables S3 and S4).

On the contrary, the antibody light chain makes fewer contacts with the RBD and around 25% of the total BSA (fig. 3A and table S3). CDRL1 and CDRL3 are part of the RBD interface in all three conformations, with only CDRL1 contributing hydrogen-bond/salt bridge interactions (tables S3 and S4). We hypothesize that the unique ability to bind both RBD -down and -up is a contributing factor to the high potency of Fab J08.

### Molecular description of the epitope-paratope interface

A commonality across the three binding conformations of J08 is residue R56 of CDRH2. In all three cases, the cryo-EM models show hydrogen bonds to the RBD backbone, and either an RBD sidechain hydrogen bond (Q493 acting as the acceptor) or salt bridge (E484) in Conformations 2 and 1, respectively (fig. 4A). In the x-ray structure of RBD-Fab J08, R56 simultaneously forms a salt bridge and hydrogen bond, respectively, with the side chains of E484 and Q493 (fig. 4A). In Conformation 3, R56 instead forms hydrogen bonds with backbone carbonyls of F490 and L492 (fig. 4A). Antibody J08 is derived from IGHV1-69*02 and is hardly mutated relative to germline (96% amino acid sequence identity with only 4 mutations observed). Unexpectedly, all 4 mutations are in CDRH2 (corresponding to Kabat numbering 55-58) (fig S3A). Relative to germline, the mutations include I56R, which is involved in key interactions as outlined above. The germline I56 residue would not be capable of side-chain hydrogen bond interactions, suggesting this mutation was selected to increase affinity and/or specificity. G55D and A57V do not directly interact with the RBD but appear to contribute to the overall stability and rigidity of CDRH2. D55 of CDRH2 forms a salt bridge with K73 of framework region 3 (FRH3), while V57, with its bulkier hydrophobic side chain relative to germline, points toward a region with several hydrophobic side chains, strengthening the interaction between beta strands of CDRH2 and FRH3 (fig. S3B). Based on the x-ray structure, N58M might also stabilize the local environment as it is part of a hydrophobic cluster that includes F486 (RBD), W47 (HC), and L96 (LC). All four of these mutations emphasize the importance of CDRH2 as the key anchoring point of J08 to the RBD.

**Fig. 4.**
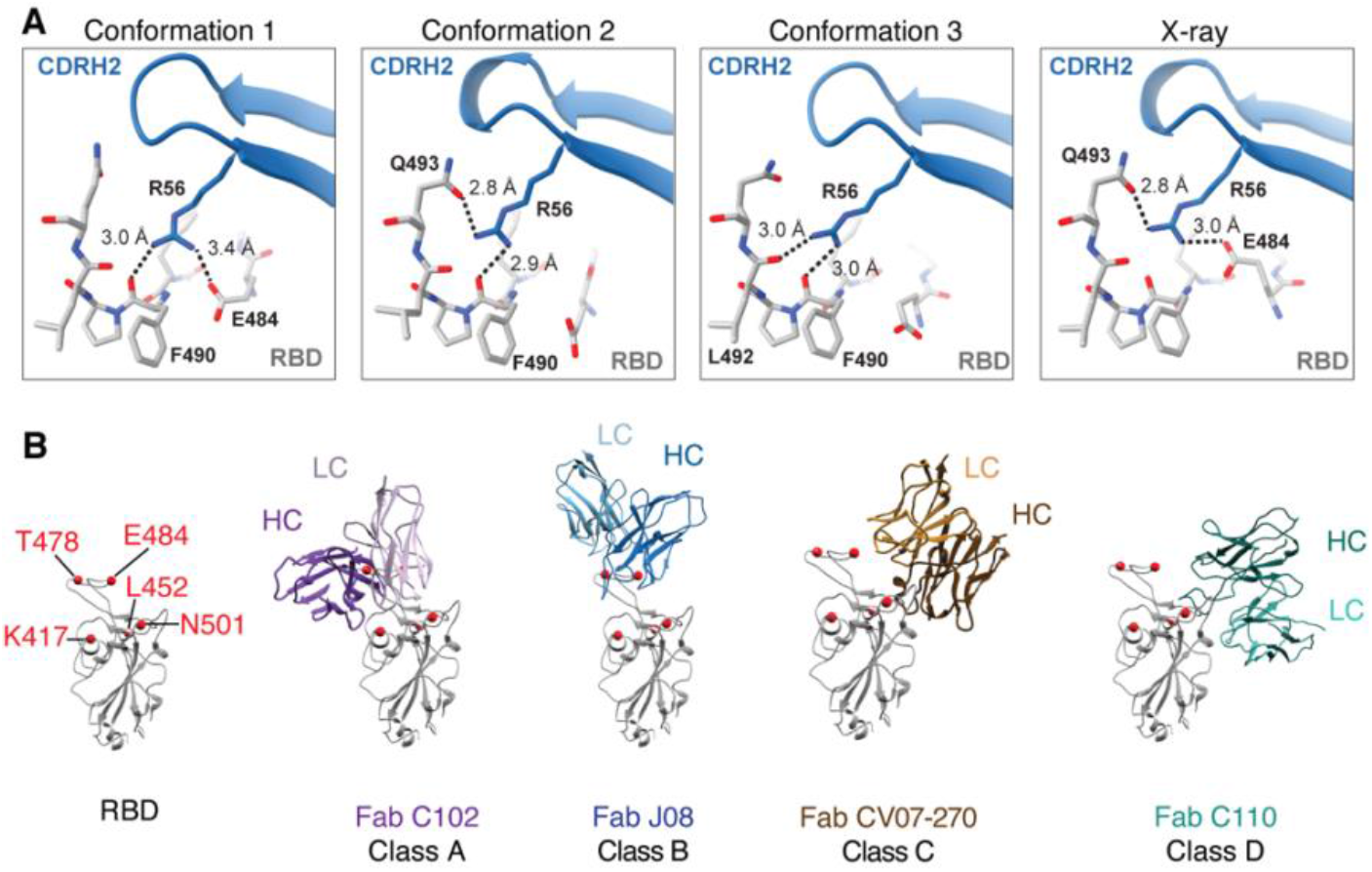
Molecular contacts and RBD epitope classification of Fab J08. (**A**) Key molecular contacts between J08 CDRH2 residue R56 and RBD are highlighted, and hydrogen bonds are represented as dashed lines with distances labeled. (**B**) Fab J08 belongs to the RBS-B class of antibodies in comparison to Fab C102, Fab CV07-270, and Fab C110 which belong to RBS-A, RBS-C, and RBS-D, respectively.

The light chain, derived from IGKV3-11, has two mutations relative to germline, one of which (L4M) might stabilize CDRL1, which is near the RBD ridge (fig. S3, A and C).

### Fab J08 does not interact with mutations found in variants of concern

Since Fab J08 binds high on the RBD ridge, it is far less susceptible to the commonly mutated residues in the receptor binding site (RBS) in the emerging variants of concern. As previously described (*29*), the RBS is subdivided into 4 epitopes, delineated as RBS-A, B, C, D. Fab J08 belongs to the RBS-B class, in comparison to other antibodies such as Fab C102 (RBS-A), Fab CV07-270 (RBS-C), and C110 (RBS-D) (fig. 4B). Many of the commonly mutated residues exist in these epitopes, thus increasing the odds of viral escape (fig. 4B; fig. S4). For example, K417 resides in RBS-A, E484 in RBS-B, L452 and E484 in RBS-C, and N501 in between RBS-A and RBS-D. Studies have shown that the binding potency and neutralization capacity of several previously isolated monoclonal antibodies is severely reduced or abrogated in the presence of one of these point mutations (*6*).

N501Y, a mutation found in the Alpha, Beta and Gamma variants, is not part of the J08 interface, explaining why neutralization against the Alpha variant is unaffected. Common to the Beta and Gamma variants are mutations at position 417 (K417N in Beta, K417T in Gamma). The E484K mutation is shared amongst the Alpha, Beta, and Gamma variants. K417 is at the interface with J08 in Conformations 1 and 3, and the x-ray structure, contributing a hydrogen bond via its side chain amine with CDRH3 in Conformations 1 and 3 (fig. S3D; tables S3 and S4). Since neutralization against Beta and Gamma variants is unaffected, we conjecture that this interaction is not critical, and becomes redundant with additional CDRH3 residues involved in the RBD interaction. E484, as mentioned, is predicted to form a salt bridge with R56 of CDRH2 in some of the conformations (table S3), so introducing a basic residue in its place might affect binding. However, neutralization suggests otherwise, and perhaps this is due to the versatility of R56 finding alternate contacts (e.g. with RBD backbone carbonyls in Conformation 3). Finally, the Delta variant has L452R and T478K mutations, and while T478 appears at the LC interface, neither residue has a measured interaction with J08 in our structures (fig. S4; table S4).

RBS-B antibodies that bind high on the RBD-ridge, share a similar angle of approach as Fab J08, and sterically block binding of hACE2, including Fab S2E12 and Fab CV07-250 (fig. 5A). In comparison to Fabs J08 and S2E12, the heavy and light chains of Fab CV07-250 are rotated approximately 90 degrees clockwise, thereby expanding the footprint on the RBD and using more light chain contacts than just primarily heavy chain contacts. In fact, this rotation and larger footprint are remarkably similar to hACE2 (fig. 5B). In decreasing order, the RBD-antibody/receptor BSA is largest for CV07-250 (943 Å^2^), followed by ACE2 (843 Å^2^), S2E12 (754 Å^2^) and J08 (676 Å^2^), based on the RBD-Fab J08 x-ray structure. Fab S2E12 is most similar to J08 based on epitope, as both antibodies make most of their contacts with the RBD via the heavy chain, and both can also neutralize variants containing the E484K/D614G and E484Q/D614G/Q779H mutations. 45% of RBD residues involved in the RBD-ACE2 interface are also part of the J08 interface, and these residues in turn account for 77% of the RBD-J08 interface (fig. S5), suggesting that escape mutations to J08 that do not also negatively impact viral fitness are rare.

**Fig. 5.**
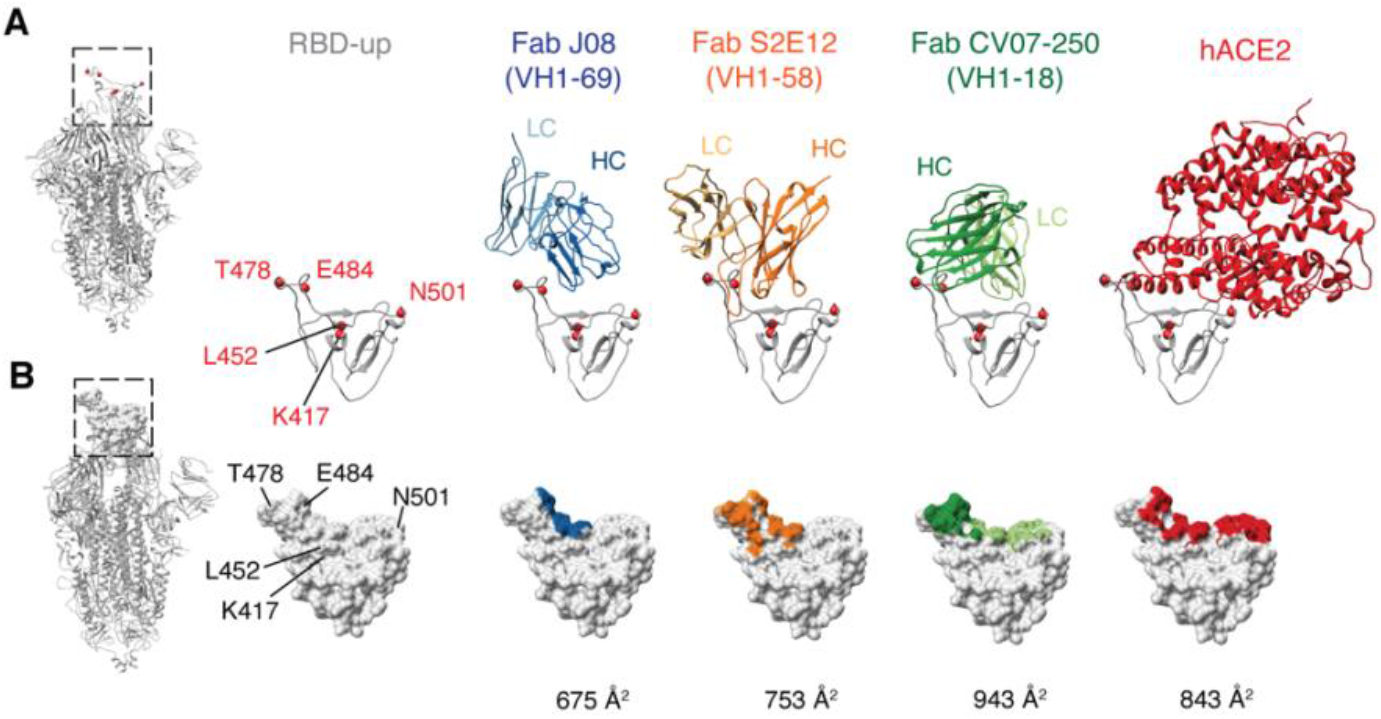
Epitope footprint comparison of Fab J08 to similar antibodies and hACE2. (**A**) Fab J08 compared to antibody Fab S2E12 and Fab CV07-250, which share similar angles of approach and bind high on the RBD ridge, thereby allosterically inhibiting hACE2 binding. Commonly mutated residues K417, L452, T478, E484, and N501 are shown as a point of reference and orientation of the RBD. (**B, C**) Surface representation of the RBD with antibody contacts colored on the surface and calculated BSA values (Å^2^) reveal that Fab J08 has the smallest footprint and therefore the least susceptible to escape mutations.

## Discussion

We used a combination of cell-based assays and structural biology to better understand the remarkable potency and resistance to variants of concern by monoclonal antibody J08. Neutralization assays using authentic virus and pseudovirus revealed that Fab J08 was able to neutralize VoCs at a low nanomolar affinity and outperform similar RBD-binding competitor antibodies, such as REGN10987/10933, S309, COV2.2196, and LY-COV016 in the low-nanomolar range. From the cryo-EM experiments, we showed that J08 can bind to multiple conformations of the spike protein with the RBD in either the -up or -down position and its smaller epitope footprint is distant from many of the common mutations found in the VOCs. E484, while on the edge of the epitope and a potential hydrogen bond or salt bridge partner, does not appear to be straddled by the antibody in a way that would create a clash in the E484K variant. mAb J08 is currently being evaluated in clinical trials to test its utility as an important intervention therapeutic for moderate-to-severe COVID. The very high potency and ability to resist escape mutations are critical properties for the next generation of SARS-CoV-2 monoclonal antibody therapeutics.

## Acknowledgements

We thank B. Anderson and H. Turner for microscope assistance, C. Bowman for computational assistance, and G.K. Hedestam and M. Corcoran for scientific discussion regarding human antibody germline genes.

## Author contributions

JLT, GO, EA, RR, and ABW designed the research, IP, NM, EP, PP and JC produced and purified the proteins. CP performed binding and ACE2 blocking assays. EA, GP, DP and TB performed BSL3 neutralization assays. LD and MC produced pseudoviruses and performed BSL2 neutralization assays. JLT performed the cryo-EM imaging and processed the data. GO built the atomic models. HL and IAW provided the RBD-Fab J08 x-ray structure. RR, and ABW coordinated the project. JLT, GO, EA, RR and ABW wrote the manuscript with input from all authors.

## Data and materials availability

Data needed to evaluate conclusions in this paper are present in both the paper itself and/or in Supplementary Materials. The cryo-EM maps have been deposited in the Electron Microscopy Data Bank (EMDB) with accession codes EMD-24876 (SARS-CoV-2-6P-Mut2), EMD-24877 (SARS-CoV-2-6P-Mut2 + Fab J08 Conformation 1), EMD-24878 (SARS-CoV-2-6P-Mut2 + Fab J08 Conformation 2), EMD-24879 (SARS-CoV-2-6P-Mut7 + Fab J08), and the atomic models deposited in the Protein Data Bank (PDB) with accession codes 7s6i (SARS-CoV-2-6P-Mut2), 7s6j (SARS-CoV-2-6P-Mut2 + Fab J08 Conformation 1), 7s6k (SARS-CoV-2-6P-Mut2 + Fab J08 Conformation 2), and 7s6l (SARS-CoV-2-6P-Mut7 + Fab J08). The coordinate and structure factor of the crystal structure of SARS-CoV-2 RBD in complex with Fab J08 has been deposited in the Protein Data Bank with accession code 7sbu.

## SUPPLEMENTARY MATERIALS

Materials and Methods

Figs. S1 to S5

Tables S1 to S4

References 30-50

