## Supplemental Material for "Structural insights of a highly potent pan-neutralizing SARS-CoV-2 human monoclonal antibody"

### Supplementary Materials for
### Structural insights of a highly potent pan-neutralizing SARS-CoV-2 human monoclonal antibody

Jonathan L. Torres[1,*], Gabriel Ozorowski[1,*], Emanuele Andreano[2], Hejun Liu[1], Jeffrey Copps[1], Giulia Piccini[3], Lorena Donnici[5], Matteo Conti[5], Cyril Planchais[6], Delphine Planas[7,8], Noemi Manganaro[2], Elisa Pantano[2], Ida Paciello[2], Piero Pileri[2], Timothée Bruel[7,8], Emanuele Montomoli[3,4,9], Hugo Mouquet[6], Olivier Schwartz[7,8], Claudia Sala[2], Raffaele De Francesco[5], Ian A. Wilson[1,10], Rino Rapuoli[2,11], Andrew B. Ward[1, §]

[1]Department of Integrative Structural and Computational Biology, The Scripps Research Institute, La Jolla, CA 92037, USA

[2]Monoclonal Antibody Discovery (MAD) Lab, Fondazione Toscana Life Sciences, Siena, Italy

[3]VisMederi S.r.l, Siena, Italy

[4]VisMederi Research S.r.l., Siena, Italy

[5]INGM, Istituto Nazionale Genetica Molecolare "Romeo ed Enrica Invernizzi", Milan, Italy.

[6]Laboratory of Humoral Immunology, Department of Immunology, Institut Pasteur, INSERM U1222, Paris, France

[7]Virus and Immunity Unit, Department of Virology, Institut Pasteur, CNRS UMR 3569, Paris, France.

[8]Vaccine Research Institute, Creteil, France

[9]Department of Molecular and Developmental Medicine, University of Siena, Siena, Italy

[10]The Skaggs Institute for Chemical Biology, The Scripps Research Institute, La Jolla, CA 92037, USA

[11]Department of Biotechnology, Chemistry and Pharmacy, University of Siena, Siena, Italy

[*]These authors contributed equally to this work

**This PDF file includes:**

Materials and Methods

Figs. S1 to S5

Tables S1 to S4

References 30-50

#### MATERIALS AND METHODS

##### Enzyme-Linked Immunoassay (ELISA)

High-binding 96-well ELISA plates (Costar, Corning) were coated overnight with 250 ng/well of purified recombinant SARS-CoV-2 proteins. After washing with 0.05% Tween 20-PBS (PBST), plates were blocked for 2 hours with 2% BSA, 1 mM EDTA, PBST (blocking buffer), washed, and incubated with purified monoclonal IgG antibodies at 10 µg/ml and 7 consecutive 1:4 dilutions in PBS. After the PBST washing, the plates were incubated with goat HRP-conjugated anti-human IgG antibodies for 1 h (Jackson ImmunoReseach, 0.8 µg/ml final in blocking buffer) and analyzed by adding 100 µl of HRP chromogenic substrate (ABTS solution, Euromedex) after the washing steps. For competition experiments of RBD-binding to ACE-2, ELISA plates were coated overnight with 250 ng/well of purified ACE-2 ectodomain. After PBST washing, plates were blocked for 2 hours with blocking buffer, washed with PBST, and incubated with purified monoclonal IgG antibodies at 10 µg/ml and 7 consecutive 1:2 dilutions in PBS in the presence of biotinylated RBD proteins at 0.5 µg/ml. After washing, the plates were incubated for 30 min with HRP-conjugated streptavidin (BD Biosciences), and analyzed by adding 100 µl of HRP chromogenic substrate (ABTS solution, Euromedex). Optical densities were measured at 405 nm ($OD_{405nm}$), and background values, assessed by incubation of PBS alone in coated wells, were subtracted. Experiments were performed using a HydroSpeed™ microplate washer and Sunrise™ microplate absorbance reader (Tecan Männedorf, Switzerland).

**SARS-CoV-2 authentic virus neutralization assay**

All SARS-CoV-2 authentic virus neutralization assays were performed in the biosafety level 3 (BSL3) laboratories at Toscana Life Sciences in Siena (Italy), Vismederi Srl, Siena (Italy) and Institute Pasteur, Paris (France). The BSL3 laboratories are approved by a Certified Biosafety Professional and inspected every year by local authorities. Two different approaches were used to evaluate the neutralization activity of J08 against SARS-CoV-2 and emerging variants and the neutralization breadth of tested antibodies. The first method is the cytopathic effect (CPE)-based neutralization assay described by Andreano and colleagues, (*13*) while the second method is a S-fuse neutralization assay previously described by Planas *et al.* (*24*). Briefly, the CPE-based neutralization assay reports on the co-incubation of mAbs with a SARS-CoV-2 viral solution containing 100 $TCID_{50}$ of virus and after 1 hour incubation at 37°C, 5% $CO_2$. The mixture was then added to the wells of a 96-well plate containing a sub-confluent Vero E6 cell monolayer. Plates were incubated for 3 days at 37°C in a humidified environment with 5% $CO_2$, then examined for CPE by means of an inverted optical microscope. As for the S-fuse neutralization assay, U2OS-ACE2 GFP1-10 or GFP 11 cells, also termed S-Fuse cells, emit fluorescence when they are productively infected by SARS-CoV-2 (*30, 31*). Cells were tested negative for mycoplasma. Cells were mixed (1:1 ratio) and plated at $8x10^3$ per well in a µClear 96-well plate (Greiner Bio-One). SARS-CoV-2 viruses were incubated with mAbs for 15 minutes at room temperature and added to S-Fuse cells. After 18 hours, cells were fixed with 2% Paraformaldehyde (PFA), washed, and stained with Hoechst Solution (1:1,000 dilution, Invitrogen). Images were acquired with an Opera Phenix high content confocal microscope (PerkinElmer). The GFP area and the number of nuclei were quantified using the Harmony software (PerkinElmer). The percentage of neutralization was calculated using the number of syncytia as value with the following formula: 100 x (1 – (value with mAb – value in “non-infected”)/(value in “no mAb” – value in “non-infected”)). We previously reported a correlation between neutralization titers obtained with the S-Fuse assay and a pseudovirus neutralization assay (*32*).

**SARS-CoV-2 virus variants for CPE-MN and S-fuse neutralization assays**

The SARS-CoV-2 viruses used to perform the CPE-MN neutralization assay were D614G (EVAg Cod: 008V-04005), B.1.1.7 (INMI GISAID accession number: EPI_ISL_736997), B.1.351 (EVAg Cod: 014V-04058), P.1 (EVAg CoD: 014V-04089) and B.1.617.2 (ID: EPI_ISL_2029113). The SARS-CoV-2 viruses used to perform the S-fuse neutralization assay were D614G, B.1.1.7, B.1.351 and B.1.617.2 and their sequences were deposited on GISAID, with the following IDs: D614G: EPI_ISL_414631; B.1.1.7: EPI_ISL_735391; B.1.1.351: EPI_ISL_964916; B.1.617.2: ID: EPI_ISL_2029113 (*25*).

**HEK293TN- hACE2 cell line generation**

An HEK293TN- hACE2 cell line was generated by lentiviral transduction of HEK293TN cells as described in Notarbartolo S. *et al.* (*33*). Briefly, HEK293TN cells were obtained from System Bioscience. Lentiviral vectors were produced following a standard procedure based on calcium phosphate co-transfection with 3$^{rd}$ generation helper and transfer plasmids. The following helper vectors were used (gifts from Didier Trono): pMD2.G/VSV-G (Addgene #12259), pRSV-Rev (Addgene #12253), pMDLg/pRRE (Addgene #12251). The transfer vector pLENTI_hACE2_HygR was obtained by cloning of hACE2 from pcDNA3.1-hACE2 (a gift from Fang Li, Addgene #145033) into pLenti-CMV-GFP-Hygro (a gift from Eric Campeau & Paul Kaufman, Addgene #17446). hACE2 cDNA was amplified by PCR and inserted under the CMV promoter of the pLenti-CMV-GFP-Hygro after GFP excision with XbaI and SalI digestion. pLENTI_hACE2_HygR is now available through Addgene (Addgene #155296). After transduction with hACE2 lentiviral vector, cells were subjected to antibiotic selection with hygromycin at 250 μg/ml. Expression of hACE2 cells was confirmed by flow cytometry staining using anti-hAce2 primary antibody (AF933, R&D

system) and rabbit anti-goat IgG secondary antibody (Alexa Fluor 647). HEK293TN-hACE2 cells were maintained in DMEM, supplemented with 10% FBS, 1% glutamine, 1% penicillin/streptomycin and 250 µg/ml Hygromicin (GIBCO) and expression of hACE2 was found to be stable after multiple passages.

**Production of SARS-CoV-2 pseudoparticles based on lentiviral vectors**

To generate SARS-CoV-2 lentiviral pseudotype particles, $5*10^6$ HEK-293TN cells were plated in a 15-cm dish in complete DMEM medium. The following day, 32 µg of reporter plasmid pLenti CMV-GFP-TAV2A-LUC Hygro, 12.5 mg of pMDLg/pRRE (Addgene #12251), 6.25 mg of pRSV-Rev (Addgene #12253) and 9 µg pcDNA3.1_ spike_del19 were co-transfected following a calcium phosphate transfection. pcDNA3.1_spike_del19 was generated by deletion of last 19aa of spike starting from pcDNA3.1-SARS2-Spike (a gift from Fang Li, Addgene plasmid # 145032) and is now available through Addgene (Addgene #155297). pLenti CMV-GFP-TAV2A-LUC Hygro was generated from pLenti CMV GFP Hygro (Addgene #17446) by addition of T2A-Luciferase by PCR cloning. 12h before transfection, the medium was replaced with complete ISCOVE. 30 h after transfection, the supernatant was collected, clarified by filtration through 0.45-µm pore-size membranes, and concentrated by centrifugation for 2h at 20,000 rpm using SW32Ti rotor. Viral pseudoparticle suspensions were aliquoted and stored at −80°C.

**SARS-CoV-2 pseudovirus neutralization assay**

Pseudovirus neutralization assays were carried out as previously described (*34*). Briefly, HEK293TN-hACE2 cells were plated at $10^4$ cells/well in white 96-well plates in complete DMEM medium. 24 hrs later, cells were infected with 0.1 MOI of SARS-CoV-2 pseudoparticles that were previously incubated with serial dilution of mAb. In particular, mAbs under test were serially diluted five-fold in PBS in order to obtain a 7-point dose-response curve (plus PBS as untreated control). Thereafter, 5 µl of each dose-response curve point was added 45 µl of medium containing SARS-

CoV-2 pseudoparticles adjusted to contain 0.1 MOI. After incubation for 1h at 37°C, 50 µl of a mAb/SARS-CoV-2 pseudoparticle mixture was added to each well and plates were incubated for 24h at 37°C. Each point was assayed in triplicate. After 24 h of incubation, cell infection was measured by a luciferase assay using the Bright-Glo™ Luciferase System (Promega) and an Infinite F200 plate reader (Tecan) to read the luminescence. Obtained RLUs were normalized to controls and dose response curve were generated by nonlinear regression curve fitting with GraphPad Prism to calculate Neutralization Dose 50 ($ND_{50}$).

**Expression and purification of SARS-CoV-2 Spike protein in the prefusion conformation**

Mutagenesis was performed on the SARS-CoV-2-6P plasmid to include S383C and D985C for the SARS-CoV-2-6P-Mut2 construct and V705C and T883C for SARS-CoV-2-6P-Mut7 construct. Expression of SARS-CoV-2-6P-Mut2 or SARS-CoV-2-6P-Mut7 S-protein was performed by incubating 0.5 mg of DNA with 1.5 mg of polyethylenimine (PEI) for 20 minutes. The mixture was placed into 1 L of HEK293F cells (Thermo Fisher), incubated for 6 days at 37°C with 8% $CO_2$ and shaken at 125 rpm. After cell harvest, the supernatant was passed over a StrepTactin XT 4FLOW column (IBA Lifesciences), washed with Buffer W (100 mM Tris-Cl pH 8, 150 mM NaCl, 1 mM EDTA), and eluted with Buffer BXT (100 mM Tris-Cl pH 8, 150 mM NaCl, 1 mM EDTA, 50 mM Biotin). The eluant was then size exclusion purified over a Superose 6 Increase-16/600 pg, 120 ml column (Cytiva). Purified trimers were buffer exchanged back into Buffer W using a 100 kDa concentrator (Amicon).

**Sample vitrification for Cryo-EM**

SARS-CoV-2-6P-Mut2 was incubated with a 3-fold molar excess of Fab J08 at room temperature for 5 minutes. The final concentration of the complex was 3 mg/ml. To aid with sample dispersal on the grid, the complex was briefly incubated with n-Dodecyl-B-D-Maltoside (DDM; final concentration 0.06 mM) and deposited on plasma-cleaned Quantifoil 1.2/1.3 4C grids. A Thermo

Fisher Vitrobot Mark IV set to 4°C, 100% humidity, 6 second wait time, and a 3 second blot time was used for the sample vitrification process. SARS-CoV-2-6P-Mut7 was incubated with a 3-fold molar excess of Fab J08 at room temperature for 30 minutes. The sample vitrification process was as described above except that the detergent was fluorinated octyl maltoside (FOM; final concentration of 0.02% w/v) and the grids were UltrAuFoil 1.2-1.3 3C.

**Cryo-EM data collection**

Datasets for both complexes were collected at 36,000x magnification on a Thermo Fisher Talos Arctica (200-keV, 1.15 Å pixel size) electron microscope with a 4k by 4k Gatan K2 Summit direct electron detector. Data collection was automated with the Leginon software (*35*) and raw micrographs were stored in the Appion database (*36*). For the SARS-CoV-2-6P-Mut2 + Fab J08 complex, a total of 2,325 micrographs were collected with a total dose of 50 e-/ Å$^2$ fractionated over 48 frames, with each frame receiving a dose rate of 5.5 electrons per pixel per second. A defocus range of -0.2 µm to -2.4 µm was used. For the SARS-CoV-2-6P-Mut7 + Fab J08 complex, 4,090 micrographs were collected with a total dose of 50 e-/ Å$^2$ but fractionated over 50 frames, with each frame receiving a dose rate of 5.2 electrons per pixel per second. In this case, a defocus range of -0.5 µm to -2.0 µm was used.

**Cryo-EM data processing, model building, and refinement**

The micrograph movie frames were aligned and dose weighted with MotionCorr2 (*37*). Aligned frames were imported into cryoSPARC v3.2 (*38*) where the CTF was estimated using Patch CTF. Particles were picked using templates (created from an initial round of 2D classification after automated picking), extracted, and subjected to multiple rounds of 2D classification for cleaning. An apo (unliganded) spike protein was imported for 3D classification (heterogeneous refinement) and the best classes were further refined. To further improve the resolution, the maps were subjected to global and local CTF refinements, and 3D variability analyses. Final refinements were performed using the non-uniform refinement feature (*39*). A summary of data collection and processing statistics can be found in table S1.

Initial models were generated by fitting Spike coordinates from PDB 6vsb and the RBD-J08 x-ray structure (see below) into the cryo-EM maps using UCSF Chimera (*40*). Several rounds of iterative manual and automated model building and relaxed refinement were performed using Coot 0.9.4 (*41*) and Rosetta (*42*). Models were validated using EMRinger (*43*) and MolProbity (*44*) as part of the Phenix software suite (*45*). Kabat numbering was applied to the antibody Fab variable light and heavy chains using the Abnum antibody numbering server (*46*). Final refinement statistics and PDB deposition codes for generated models can be found in table S1. Buried surface area calculations and distance measurements were performed using PDBePISA (*47*).

**Crystallization and X-ray structure determination**

The J08 Fab complexed with SARS-CoV-2 RBD was formed by mixing each of the protein components in an equimolar ratio and incubating overnight at 4°C. 384 conditions of the JCSG Core Suite (Qiagen) were used for setting-up trays for the complex (6 mg/mL) on robotic CrystalMation system (Rigaku) at Scripps Research. Crystallization trials were set-up by the vapor diffusion method in sitting drops containing 0.1 μl of protein complex and 0.1 μl of reservoir solution. Crystals appeared on day 3, were harvested on day 7, pre-equilibrated in cryoprotectant containing 10% ethylene glycol, and then flash cooled and stored in liquid nitrogen until data collection. Diffraction data were collected at cryogenic temperature (100 K) at beamline 12-1 of the Stanford Synchrotron Radiation Lightsource (SSRL) and processed with HKL2000 (*48*). Diffraction data were collected from crystals grown in drops containing 17% (w/v) PEG 4000, 15% (v/v) Glycerol, 8.5% (v/v) Isopropanol, 0.085 M Sodium HEPES pH 7.5. The X-ray structures were solved by molecular replacement (MR) using PHASER (*49*) with MR models for the RBD and Fab from PDB 7JMW (*50*). Iterative model building and refinement were carried out in COOT (*41*) and

PHENIX (*45*), respectively. x-ray data collection and structural refinement statistics can be found in table S2.

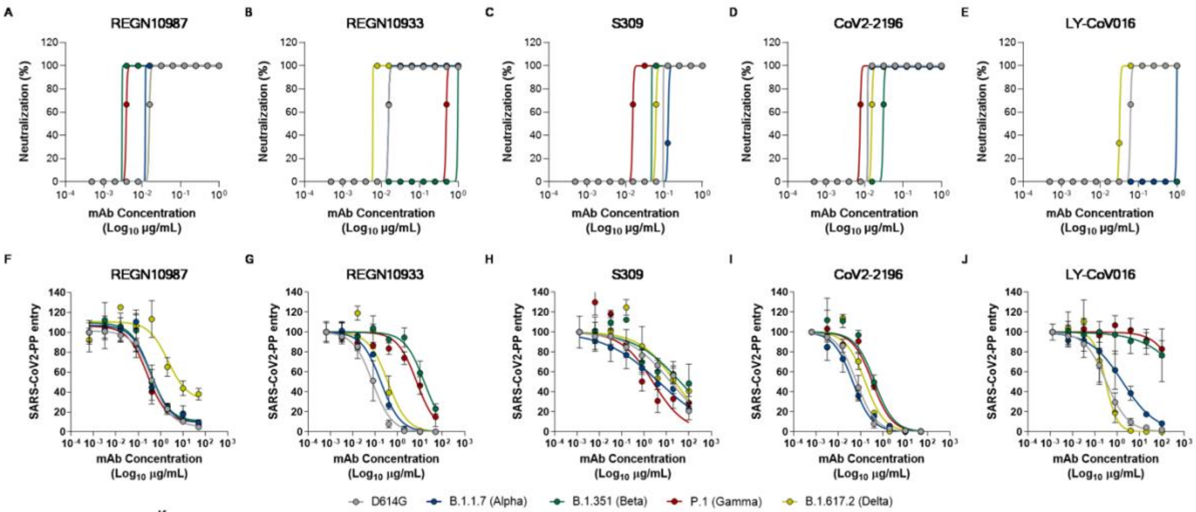


K

| mAb ID | CPE-MN ($IC_{100}$ ng/mL) | | | | | Pseudovirs ($IC_{50}$ ng/mL) | | | | |
|---|---|---|---|---|---|---|---|---|---|---|
| | D614G | B.1.1.7 (Alpha) | B.1.351 (Beta) | P.1 (Gamma) | B.1.617.2 (Delta) | D614G | B.1.1.7 (Alpha) | B.1.351 (Beta) | P.1 (Gamma) | B.1.617.2 (Delta) |
| REGN10987 | 24.6 | 19.5 | 4.9 | 3.1 | 19.7 | 41.0 | 37.0 | 35.0 | 25.0 | 2,154 |
| REGN10933 | 24.6 | 24.6 | 1,574 | 787.4 | 7.8 | 75.0 | 179 | 15,460 | 7,120 | 365 |
| S309 | 156.2 | 248.0 | 78.1 | 24.6 | 78.7 | 15,000 | 4,600 | 41,000 | 2,500 | 24,000 |
| CoV2-2196 | 19.5 | 19.5 | 49.2 | 12.3 | 19.7 | 66.0 | 45.0 | 425.0 | 335.0 | 161.0 |
| LY-CoV016 | 124.0 | 1,574 | Not-neutralizing | Not-neutralizing | 49.6 | 326.0 | 1,744 | Not-neutralizing | Not-neutralizing | 305.0 |

**Fig. S1. Neutralization activity of competitor mAbs**. (**A-E**) Graphs show the CPE-MN neutralization activity against SARS-CoV-2 D614G, B.1.1.7, B.1.351, P.1, and B.1.617.2 for REGN10987 (**A**), REGN10933 (**B**), S309 (**C**), CoV2-2196 (**D**) and LY-CoV016 (**E**). (**F-J**) Graphs show the neutralization activity against SARS-CoV-2 D614G, B.1.1.7, B.1.351, P.1, and B.1.617.2 pseudoviruses for REGN10987 (**F**), REGN10933 (**G**), S309 (**H**), CoV2-2196 (**I**) and LY-CoV016 (**J**). (**K**) The table summarizes the $IC_{100}$ and $IC_{50}$ results obtained for all neutralization assays.

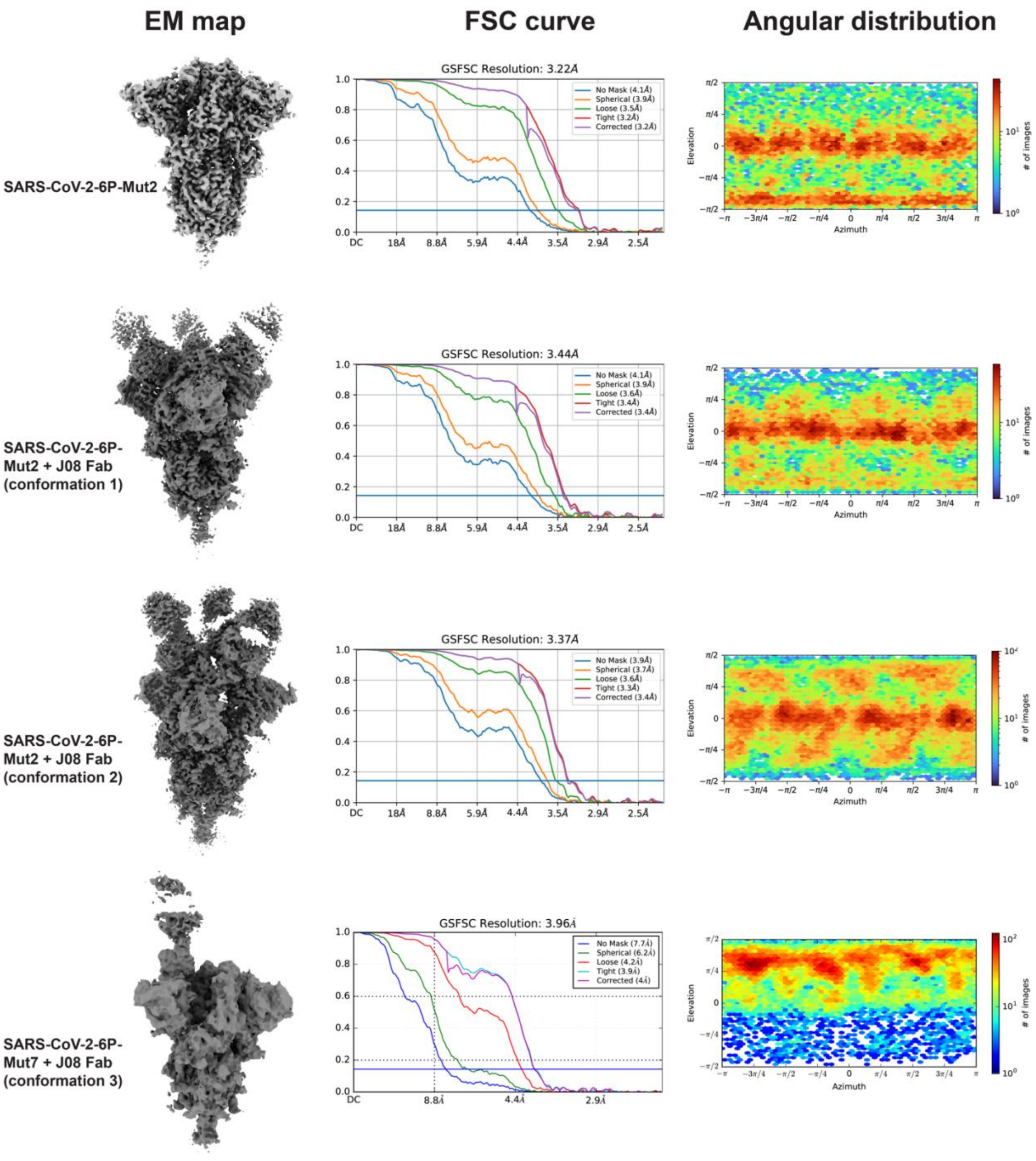


**Fig. S2. Cryo-EM resolution estimates and angular sampling.** EM map (*left*), Fourier Shell Correlation (FSC) resolution curves (*middle*) and angular distribution plot (*right*) of the four cryo-EM reconstructions.

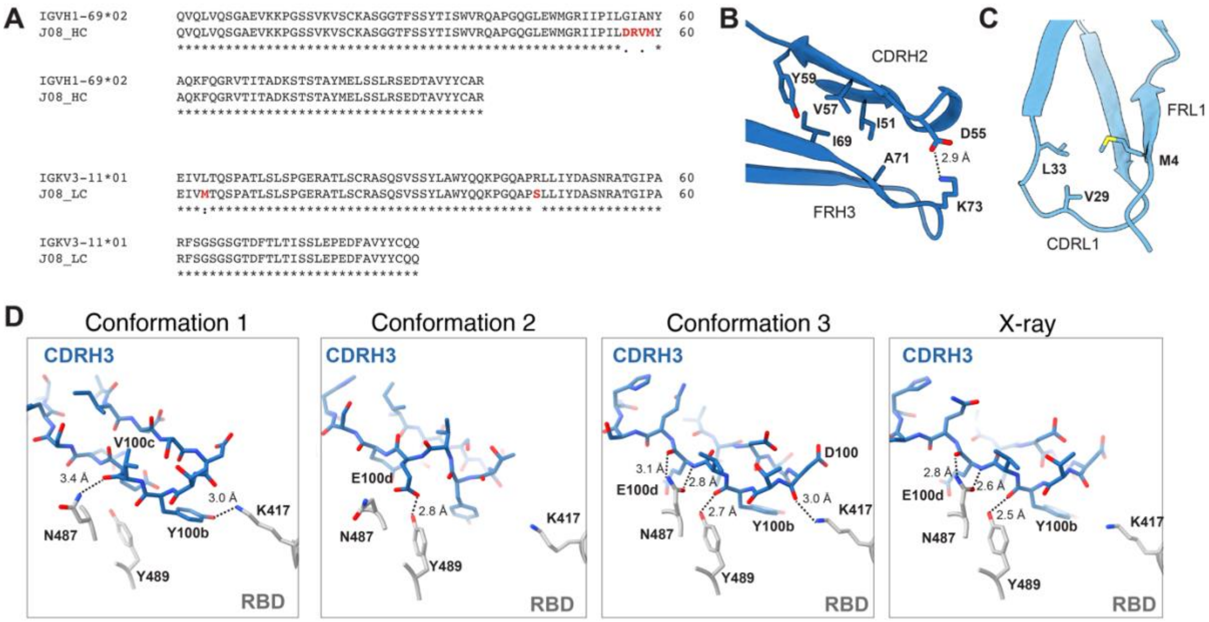


**Fig. S3. Key J08 mutations relative to the predicted germline sequence.** (**A**) Sequence alignment of J08 heavy and light chains with their respective predicted germline V genes reveals 4 mutations in CDRH2, and 2 mutations in the framework regions of the light chain. (**B**) The heavy chain G55D and A57V mutations stabilize the interaction between CDRH2 and FRH3. (**C**) An L4M mutation in the J08 light chain might stabilize CDRL1 through additional hydrophobic interactions. (**D**) Interactions between RBD and J08 CDRH3 in all four J08-bound structures. Predicted hydrogen bonds represented as dotted lines with distances labeled.

A

| Origin | UK | South Africa | Brazil | California | California | New York | New York | India | India |
|---|---|---|---|---|---|---|---|---|---|
| Pango Lineage | B1.1.7 | B.1.351 | P.1 | B.1.427 | B.1.429 | B.1.525 | B.1.526 | B.1.617 | B.1.617.2 |
| WHO Label | Alpha | Beta | Gamma | Epsilon | Epsilon | Eta | Iota | Delta | Delta |
| NTD | 69-70 del<br>Y144 del | D80A<br>D215G | L18F<br>T20N<br>P26S<br>D138Y<br>R190S | | S13I<br>W152C | Q52R<br>69-70 del<br>Y144 del | L5F<br>T95I<br>D253G | G142D<br>E154K | T19R<br>G142D<br>156 del<br>157 del<br>R158G |
| RBD | E484K<br>N501Y | K417N<br>E484K<br>N501Y | K417T<br>E484K<br>N501Y | L452R | L452R | E484K | S477N<br>E484K | E484Q<br>L452R | L452R<br>T478K |
| S1+S2* | A570D<br>D614G<br>P681H<br>T716I<br>S982A<br>D1118H | D614G<br>A701V | D614G<br>H655Y<br>T1027I | D614G | D614G | D614G<br>Q677H<br>F888L | D614G<br>A701V | D614G<br>P681R<br>Q1071H | D614G<br>P681R<br>D950N |

B

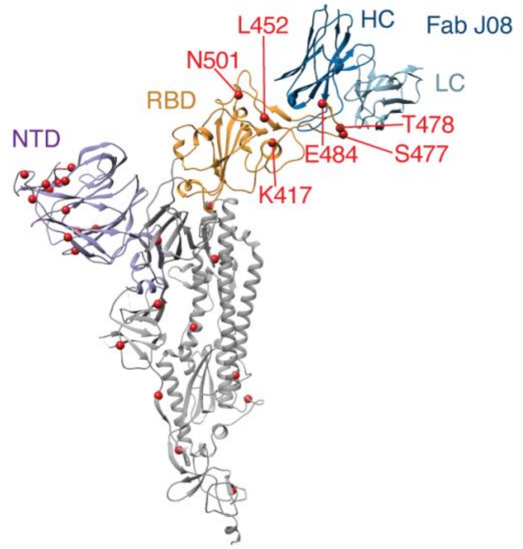


**Fig. S4**. **J08 binds away from most VOC mutations.** (**A**) Summary table and (**B**) protomer of SARS-CoV-2-Mut2 + Fab J08. Mutations listed in the summary table residing in the NTD and RBD are colored in purple and orange. Mutations not residing in the NTD or RBD are labelled as S1+S2* and colored in gray. Mutations are represented as red spheres in the protomer model.

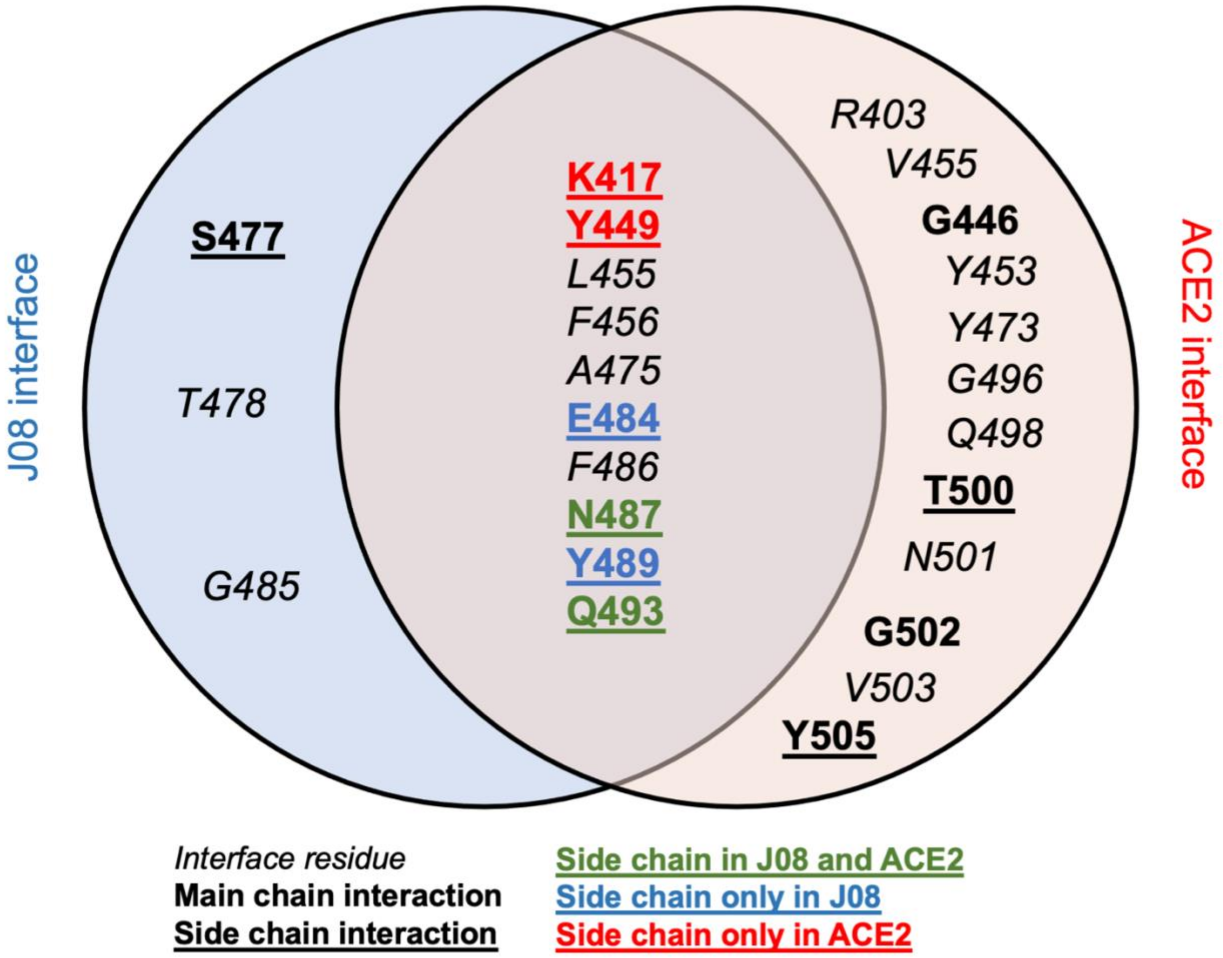


**Fig. S5**. **Venn diagram depicting shared RBD interface contacts between J08 and ACE2**. Font style and color represents whether molecular interactions between ACE2 or J08 and RBD involve side chain or backbone atoms.

#### Table S1. Cryo-EM data collection, refinement and model building statistics

| Map | SARS-CoV-2-6P-Mut2 S | J08 Fab + SARS-CoV-2-6P-Mut2 S (Conformation 1) | J08 Fab + SARS-CoV-2-6P-Mut2 S (Conformation 2) | J08 Fab + SARS-CoV-6P-Mut7 S (Conformation 3) |
|---|---|---|---|---|
| EMDB | **EMD-24876** | **EMD-24877** | **EMD-24878** | **EMD-24879** |
| **Data collection** | | | | |
| Microscope | FEI Talos Arctica | | | FEI Talos Arctica |
| Voltage (kV) | 200 | | | 200 |
| Detector | Gatan K2 Summit | | | Gatan K2 Summit |
| Recording mode | Counting | | | Counting |
| Nominal magnification | 36,000 | | | 36,000 |
| Movie micrograph pixel size (Å) | 1.15 | | | 1.15 |
| Dose rate ($e^-$/[(camera pixel)*s]) | 5.5 | | | 5.2 |
| Number of frames per movie micrograph | 48 | | | 50 |
| Frame exposure time (ms) | 250 | | | 250 |
| Movie micrograph exposure time (s) | 12.0 | | | 12.5 |
| Total dose ($e^-$/Å$^2$) | 50 | | | 50 |
| Defocus range (µm) | -0.2 to -2.4 | | | -0.5 to -2.0 |
| **EM data processing** | | | | |
| Number of movie micrographs | 2,325 | 2,325 | 2,325 | 4,090 |
| Number of molecular projection images in map | 27,831 | 32,769 | 52,678 | 43,511 |
| Symmetry | C3 | C3 | C3 | C1 |
| Map resolution (FSC 0.143; Å) | 3.2 | 3.4 | 3.4 | 4.0 |
| Map sharpening B-factor (Å$^2$) | -82.1 | -84.3 | -91.6 | -79.3 |
| **Structure building and validation** | | | | |
| *Number of atoms in deposited model* | | | | |
| SARS-CoV-2 S protein | 25,848 | 25,176 | 25,116 | 22,633 |
| Glycans | 924 | 462 | 630 | 420 |
| J08 Fv | N/A | 5,157 | 5,157 | 1,702 |
| MolProbity score | 0.82 | 0.98 | 0.95 | 0.73 |
| Clash score | 0.34 | 0.61 | 0.74 | 0.55 |
| Map correlation coefficient | 0.85 | 0.82 | 0.82 | 0.73 |
| EMRinger score | 4.14 | 2.74 | 3.18 | 1.89 |
| *RMSD from ideal* | | | | |
| Bond length (Å) | 0.02 | 0.02 | 0.02 | 0.02 |
| Bond angles (˚) | 1.72 | 1.80 | 1.79 | 1.73 |
| *Ramachandran plot* | | | | |
| Favored (%) | 96.80 | 95.95 | 96.64 | 97.83 |
| Allowed (%) | 3.20 | 4.05 | 3.36 | 2.17 |
| Outliers (%) | 0.00 | 0.00 | 0.00 | 0.00 |
| Side chain rotamer outliers (%) | 0.10 | 0.18 | 0.00 | 0.04 |
| PDB | **7s6i** | **7s6j** | **7s6k** | **7s6l** |

#### Table S2. Crystallographic data collection and refinement statistics

| | |
|---|---|
| **Data collection** | |
| Beamline | SSRL 12-1 |
| Wavelength (Å) | 0.97946 |
| Space group | $P\ 2_1\ 2_1\ 2_1$ |
| Unit cell parameters | |
| a, b, c (Å) | 55.5, 103.2, 123.0 |
| α, β, γ (°) | 90, 90, 90 |
| Resolution (Å) [a] | 50.0–2.54 (2.58–2.54) |
| Unique reflections [a] | 23,706 (1,144) |
| Redundancy [a] | 7.9 (7.7) |
| Completeness (%) [a] | 99.7 (97.4) |
| $<I/\sigma_I>$ [a] | 14.5 (3.7) |
| $R_{sym}$[b] (%) [a] | 12.8 (50.1) |
| $R_{pim}$[b] (%) [a] | 4.9 (18.6) |
| $CC_{1/2}$[c] (%) [a] | 98.5 (89.9) |
| **Refinement statistics** | |
| Resolution (Å) | 40.8–2.54 |
| Reflections (work) | 22,417 |
| Reflections (test) | 1,166 |
| $R_{cryst}$[d] / $R_{free}$[e] (%) | 22.0/26.1 |
| No. of atoms | 4,904 |
| Macromolecules | 4,750 |
| Glycans | 14 |
| Solvent | 140 |
| Average *B*-value (Å$^2$) | 37 |
| Macromolecules | 37 |
| Fab | 35 |
| RBD | 41 |
| Glycans | 59 |
| Solvent | 34 |
| Wilson *B*-value (Å$^2$) | 36 |
| **RMSD from ideal geometry** | |
| Bond length (Å) | 0.002 |
| Bond angle (º) | 0.454 |
| **Ramachandran statistics (%)**[f] | |
| Favored | 97.4 |
| Outliers | 0.0 |
| **PDB code** | 7sbu |

[a] Numbers in parentheses refer to the highest resolution shell.
[b] $R_{sym} = \Sigma_{hkl} \Sigma_i | I_{hkl,i} - <I_{hkl}> | / \Sigma_{hkl} \Sigma_i I_{hkl,i}$ and $R_{pim} = \Sigma_{hkl} (1/(n-1))^{1/2} \Sigma_i | I_{hkl,i} - <I_{hkl}> | / \Sigma_{hkl} \Sigma_i I_{hkl,i}$, where $I_{hkl,i}$ is the scaled intensity of the i$^{th}$ measurement of reflection h, k, l, $<I_{hkl}>$ is the average intensity for that reflection, and *n* is the redundancy.
[c] $CC_{1/2}$ = Pearson correlation coefficient between two random half datasets.
[d] $R_{cryst} = \Sigma_{hkl} | F_o - F_c | / \Sigma_{hkl} | F_o |$ x 100, where $F_o$ and $F_c$ are the observed and calculated structure factors, respectively.
[e] $R_{free}$ was calculated as for $R_{cryst}$, but on a test set comprising 5% of the data excluded from refinement.
[f] From MolProbity (*44*).

**Table S3. List of interactions between J08 and SARS-CoV-2 Spike.** Calculated using PDBePISA (*47*) using a cutoff distance of 3.4 Å.

***Conformation 1***

| # | Antibody region | Ab residue[atom] | RBD residue(atom) | Distance (Å) |
|---|---|---|---|---|
| **1** | CDRH2 | R50[NH2] | Y489[OH] | 3.3 |
| **2** | CDRH2 | I53[O] | Q493[NE2] | 3.1 |
| **3** | CDRH2 | R56[NH1] | E484[OE1] | 3.4 |
| **4** | CDRH2 | R56[NH2] | F490[O] | 3 |
| **5** | CDRH3 | Y100b[OH] | K417[NZ] | 3 |
| **6** | CDRH3 | V100c[O] | N487[ND2] | 3.4 |
| **7** | CDRL1 | S30[OG] | S477[OG] | 3 |
| **8** | CDRL1 | Y32[OH] | N487[OD1] | 2.7 |

***Conformation 2***

| # | Antibody region | Ab residue[atom] | RBD residue(atom) | Distance (Å) |
|---|---|---|---|---|
| **1** | FR-H1 | G27[N] | T500[O]* | 3 |
| **2** | CDRH1 | Y32[OH] | P499[O]* | 2.8 |
| **3** | CDRH2 | R50[NH1] | F486[O] | 2.9 |
| **4** | CDRH2 | L54[O] | Q493[NE2] | 2.9 |
| **5** | CDRH2 | R56[NH1] | F490[O] | 2.9 |
| **6** | CDRH2 | R56[[NH2] | Q493[OE1] | 2.8 |
| **7** | CDRH3 | A96[O] | N440[ND2]* | 3.2 |
| **8** | CDRH3 | E100d[OE2] | Y489[OH] | 2.8 |
| **9** | CDRL1 | S30[OG] | S477[OG] | 3.3 |
| **10** | CDRL1 | Y32[OH] | S477[N] | 3.1 |

***Conformation 3***

| # | Antibody region | Ab residue[atom] | RBD residue(atom) | Distance (Å) |
|---|---|---|---|---|
| **1** | CDRH2 | R56[NH1] | F490[O] | 3 |
| **2** | CDRH2 | R56[NH2] | L492[O] | 3 |
| **3** | CDRH3 | D100[O] | K417[NZ] | 3 |
| **4** | CDRH3 | Y100b[O] | Y489[OH] | 2.7 |
| **5** | CDRH3 | E100d[N] | N487[OD1] | 2.8 |
| **6** | CDRH3 | E100d[O] | N487[ND2] | 3.1 |

***X-ray***

| # | Antibody region | Ab residue[atom] | RBD residue(atom) | Distance (Å) |
|---|---|---|---|---|
| **1** | CDRH2 | L54[O] | Q493[NE2] | 3.1 |
| **2** | CDRH2 | R56[NH1] | E484[OE1] | 3 |
| **3** | CDRH2 | R56[NH2] | Q493[OE1] | 2.8 |
| **4** | CDRH3 | Y100b[O] | Y489[OH] | 2.5 |

| | | | | |
|---|---|---|---|---|
| **5** | CDRH3 | E100d[N] | N487[OD1] | 2.6 |
| **6** | CDRH3 | E100d[O] | N487[ND2] | 2.8 |
| **7** | CDRL1 | S30[OG] | S477[O] | 3.3 |
| **8** | CDRL1 | Y32[OH] | S477[OG] | 3 |

*Interaction involving adjacent protomer

**Table S4. List of interface residues between J08 and SARS-CoV-2 Spike.** Defined as contributing >5 Å$^2$ buried surface area as calculated using PDBePISA (*47*).

| ***Heavy chain*** | **Conformation1** | **Conformation2** | **Conformation3** | **X-ray** |
|---|---|---|---|---|
| **V2** | | RBD* | | |
| **G26** | | RBD* | | |
| **G27** | | RBD* | | |
| **S31** | | RBD* | | |
| **Y32** | | RBD* | | |
| **R50** | RBD | RBD | RBD | RBD |
| **I52** | | RBD | RBD | RBD |
| **I53** | RBD | | | |
| **L54** | RBD | RBD | RBD | RBD |
| **D55** | RBD | RBD | RBD | RBD |
| **R56** | RBD | RBD | RBD | RBD |
| **M58** | RBD | RBD | RBD | RBD |
| **R95** | | RBD | RBD | RBD |
| **A96** | | RBD* | | |
| **I97** | | RBD* | | |
| **D100** | RBD | RBD* | RBD | RBD |
| **T100a** | RBD | | RBD | RBD |
| **Y100b** | RBD | RBD | RBD | RBD |
| **V100c** | RBD | RBD | RBD | RBD |
| **E100d** | RBD | RBD | RBD | RBD |
| **Q100e** | RBD | RBD | | |
| **S100f** | | RBD | RBD | RBD |
| **Y102** | | RBD* | | |

| ***Light chain*** | **Conformation1** | **Conformation2** | **Conformation3** | **X-ray** |
|---|---|---|---|---|
| **S28** | | | | RBD |
| **V29** | | | RBD | RBD |
| **S30** | RBD | RBD | RBD | RBD |
| **Y32** | RBD | RBD | RBD | RBD |
| **T56** | | RBD* | | |
| **P91** | RBD | RBD | RBD | RBD |
| **L92** | RBD | RBD | | |
| **L96** | | | | RBD |

| *RBD* | Conformation1 | Conformation2 | Conformation3 | X-ray |
|---|---|---|---|---|
| **K417** | HC | | HC | HC |
| **N439** | | HC# | | |
| **N440** | | HC# | | |
| **V445** | | HC# | | |
| **Y449** | HC | | HC | HC |
| **L455** | HC | HC | HC | HC |
| **F456** | HC | HC | HC | HC |
| **Y473** | HC | | HC | |
| **A475** | HC | HC | HC | HC |
| **G476** | | | HC | LC |
| **S477** | LC | LC | LC | LC |
| **T478** | LC | LC | LC | LC |
| **V483** | | HC | | |
| **E484** | HC | | HC | HC |
| **G485** | HC | HC | HC | HC |
| **F486** | HC+LC | HC+LC | HC+LC | HC+LC |
| **N487** | HC+LC | HC | HC+LC | HC+LC |
| **Y489** | HC | HC | HC | HC |
| **F490** | HC | HC | HC | HC |
| **L492** | | | HC | |
| **Q493** | HC | HC | HC | HC |
| **P499** | | HC# | | |
| **T500** | | HC# | | |
| **V503** | | HC# | | |
| **Q506** | | HC# | | |

*Adjacent protomer RBD

#Contacts involving secondary (adjacent protomer) RBD
